# *O*-GlcNAc transferase plays a non-catalytic role in *C. elegans* male fertility

**DOI:** 10.1101/2022.05.26.493566

**Authors:** Daniel Konzman, Tetsunari Fukushige, Mesgana Dagnachew, Michael Krause, John A. Hanover

**Affiliations:** Laboratory of Cell and Molecular Biology, National Institute of Diabetes and Digestive and Kidney Diseases, National Institutes of Health, Bethesda, Maryland 20892; Johns Hopkins University Department of Biology, 3400 N. Charles Street, Baltimore, MD, 21218, USA; Laboratory of Molecular Biology, National Institute of Diabetes and Digestive and Kidney Diseases, National Institutes of Health, Bethesda, Maryland 20892

## Abstract

Animal behavior is influenced by the competing drives to maintain energy and to reproduce. The balance between these evolutionary pressures and how nutrient signaling pathways intersect with mating drive remains unclear. The nutrient sensor *O*-GlcNAc transferase, which post-translationally modifies intracellular proteins with a single monosaccharide, is responsive to cellular nutrient status and regulates diverse biological processes. Though essential in most metazoans, *O*-GlcNAc transferase (*ogt-1*) is dispensable in *Caenorhabditis elegans*, allowing genetic analysis of its physiological roles. Compared to control, *ogt-1* males have a four-fold reduction in mean offspring, with nearly two thirds producing zero progeny. Interestingly, we found that isolated *ogt-1* males are less likely to engage in mate-searching, and they initiate mating less often when exposed to mates. In addition, *ogt-1* males which do initiate mating are less likely to continue with subsequent steps in the mating process, resulting in fewer successful sperm transfers. Lowering barriers to mating such as immobilizing mates or allowing more mating time significantly improves *ogt-1* male mating. Surprisingly, we found high fertility levels for *ogt-1* mutant males with hypodermal expression of wild-type *ogt-1* and by *ogt-1* harboring mutations that prevent the transfer of *O*-GlcNAc by OGT-1. This suggests OGT-1 serves a non-catalytic function in the hypodermis impacting the male mating drive. This study builds upon research on the nutrient sensor *O*- GlcNAc transferase and demonstrates a role it plays in the interplay between the evolutionary drives for reproduction and survival.

**Author Summary:** Animals must make decisions on whether to engage in reproduction or conserve energy. These decisions must take into account the energy available to the animal, therefore making the nutrient sensing enzyme OGT of particular interest. In response to nutrient levels in the cell, OGT transfers the GlcNAc sugar onto proteins to regulate their function. OGT is implicated in a number of human diseases including diabetes, cancer, and X-linked intellectual disability. By deleting the gene encoding OGT in the nematode *C. elegans*, we show OGT is required for male fertility. We assessed the behavior of these mutant male worms and found they have a reduced mating drive. Surprisingly, restoring OGT specifically in the hypodermis was able to raise male fertility and mating drive back to normal levels. In addition, missense mutations in the OGT catalytic domain which prevent the enzyme from transferring GlcNAc do not negatively impact fertility, suggesting a different function of OGT is important in this process. Our study demonstrates that OGT is important in critical behavioral decisions and that further investigation in *C. elegans* may help reveal new functions of the enzyme.

## Introduction

The survival of an organism depends on its ability to respond to the environment around it. To maintain life and health, organisms must adequately respond to changes in factors such as temperature, toxins, and nutrition. Critically, reproduction is impacted by many environmental factors including diet, xenobiotics, and stress [1]. Two key evolutionary pressures, survival and reproduction, are both impacted by the environment and can conflict with each other. Survival is supported by the conservation of energy, acquisition of nutrients, and the avoidance of risk. Conversely, reproduction entails risks and requires extensive energy use through gametogenesis, mating, and the raising of young. The behavioral prioritization between these two drives is complex and is particularly impacted by nutrient status [2-4].

With its quick generation time and genetic amenability, the nematode *Caenorhabditis elegans* (*C. elegans*) provides a good model to investigate reproduction and behavior. While the self-fertile hermaphrodites have no need to mate in order to reproduce, males must seek out mates and go through a complex series of behaviors to successfully sire progeny [5]. After successfully locating a mate, the male must make contact with its tail, start backwards locomotion, locate the vulva, and transfer sperm. If the male starts on the side without the vulva or doesn’t detect it at first pass, it will need to execute a tight turn of its tail to switch sides and continue searching. The mating drive of males is impacted by nutrition and related pathways such as the insulin-like signaling pathway [3], making the nutrient-sensing enzyme *O-*GlcNAc transferase (OGT) a potential regulator of mating.

OGT transfers *O-*GlcNAc to serine and threonine residues of its protein targets to regulate many processes within cells. This post-translational modification is added to thousands of nucleocytoplasmic proteins, and can be dynamically removed by the *O-*GlcNAcase (OGA) [6, 7]. *O-*GlcNAc modification of proteins can modify their activity, subcellular localization, and interactions with other proteins [7, 8]. *O-* GlcNAc has been shown to regulate a diverse array of cellular processes including signaling, stress response, and gene expression [9-11]. The biosynthesis of UDP-GlcNAc, the metabolite required for *O-* GlcNAc transfer, requires inputs from many different metabolic pathways resulting in its concentration being dependent upon the overall cellular nutrient status [10, 11]. Deregulation of this process is a feature of human disorders including diabetes, cancer, and Alzheimer’s disease [8, 12].

OGT has functions other than transferring *O-*GlcNAc including HCF-1 cleavage [13], and non-catalytic roles such as protein-protein interactions, with each of these three activities being critical for growth of mammalian cells [14]. The importance of OGT is underscored by recent findings that variants in OGT cause a form of X-Linked intellectual disability (XLID) known as OGT-XLID [15, 16]. This syndrome is characterized by low IQ, developmental delay, dysmorphia, and hypotonia, among other symptoms [16]. The variants that cause OGT-XLID predominantly occur in the tetratricopeptide repeat (TPR) domain rather than the catalytic domain, and most do not ablate the enzyme’s ability to transfer *O-* GlcNAc [16]. Thus, a better understanding of OGT biology, and both its catalytic and non-catalytic functions, may provide insights into this disorder and others affected by OGT disfunction.

Though OGT is essential in most metazoans [17], loss of *O-*GlcNAc transferase (*ogt-1*) in *C. elegans* is viable [18]. As the only model organism that tolerates loss of OGT, *C. elegans* is uniquely suited for genetic analysis of the biological roles of this enzyme. *ogt-1* mutant worms have proven to be a powerful model of insulin resistance [11, 18, 19], stress [20-24], and neurological disorders [17, 25-28]. Highlighting the gene’s importance in metabolic regulation, *ogt-1* deletion causes altered macronutrient storage, including a three-fold reduction in lipid stores and three-fold increase in glycogen and trehalose stores [18]. These animals also have altered entry into diapause states when starved, being more likely to form dauers [18, 29], and less likely to enter adult reproductive diapause [30]. Developmental changes have also been noted, including a decreased lifespan [21, 31] and increased susceptibility to transdifferentiation [32, 33]. Further, *ogt-1* mutant worms have been used to uncover the role of OGT in such behaviors as hypersensitivity to touch and altered habituation to repeated stimuli [34]. While *ogt-1* hermaphrodites have only a slight reduction in brood size in normal husbandry conditions [20, 23, 31, 35], we noticed males produce very few offspring in crosses, despite no obvious uncoordinated movement phenotypes. Here we detail the *ogt-1* male fertility defect and provide evidence this phenotype relies on a non-catalytic role of OGT-1 in the hypodermis that results in behavioral prioritization away from mate-seeking behavior.

## Results

### *ogt-1* is required for male fertility

We set out to determine the impact that *ogt-1* deletion has on male fertility. This was assessed by mating them with feminized worms and counting their progeny. *fem-1(hc17)* worms raised at 25°C produce no sperm and thus have no self-progeny, but produce offspring after mating with a fertile male [36]. In this experiment, the two control lines (N2 and *him-5*) have similarly high progeny counts (Fig 1A), and thus *him-5* was used as the genetic background “wild-type” throughout this study as the “high incidence of males” phenotype (Him) provides adequate numbers of males to conduct experiments [37]. We tested multiple alleles of *ogt-1* in the *him-5* background for male fertility and compared them with controls. *fem-1* worms mated to males with deletions in *ogt-1* had severely reduced 1-day broods compared with both *him-5* and N2 males. Three independent *ogt-1* alleles had the male fertility defect: our CRISPR deletion of the entire *ogt-1* coding region (*jah01*) [30], and two smaller deletions within the coding sequence both expected to produce no functional protein (*ok430* [18] and *ok1474* [20]) (Fig 1A). As the phenotype was strongest in the CRISPR deletion, we used this line throughout the study and refer to the allele as “*ogt-1*” unless noted otherwise. We found that the reduction of *ogt-1* male brood size was about 65% to 80% (Fig 1A). While the progeny count from *ogt-1* males was often very low or zero, all three alleles had a wide range of values (Fig 1A). We also assessed the fertility of males harboring mutations in *oga-1*, the gene encoding the enzyme that removes *O-*GlcNAc from proteins. Two *oga-1* alleles (CRISPR deletion of the entire *oga-1* coding region *av82* [30], and *ok1207* [19]) did not appear to affect male fertility, as their progeny counts were similar to positive controls (Fig 1A). Analysis of the *ogt-1(jah01);oga-1(av82)* double CRISPR deletion line showed low fertility similar to *ogt-1(jah01)* alone (Fig 1A).

**Fig 1.**
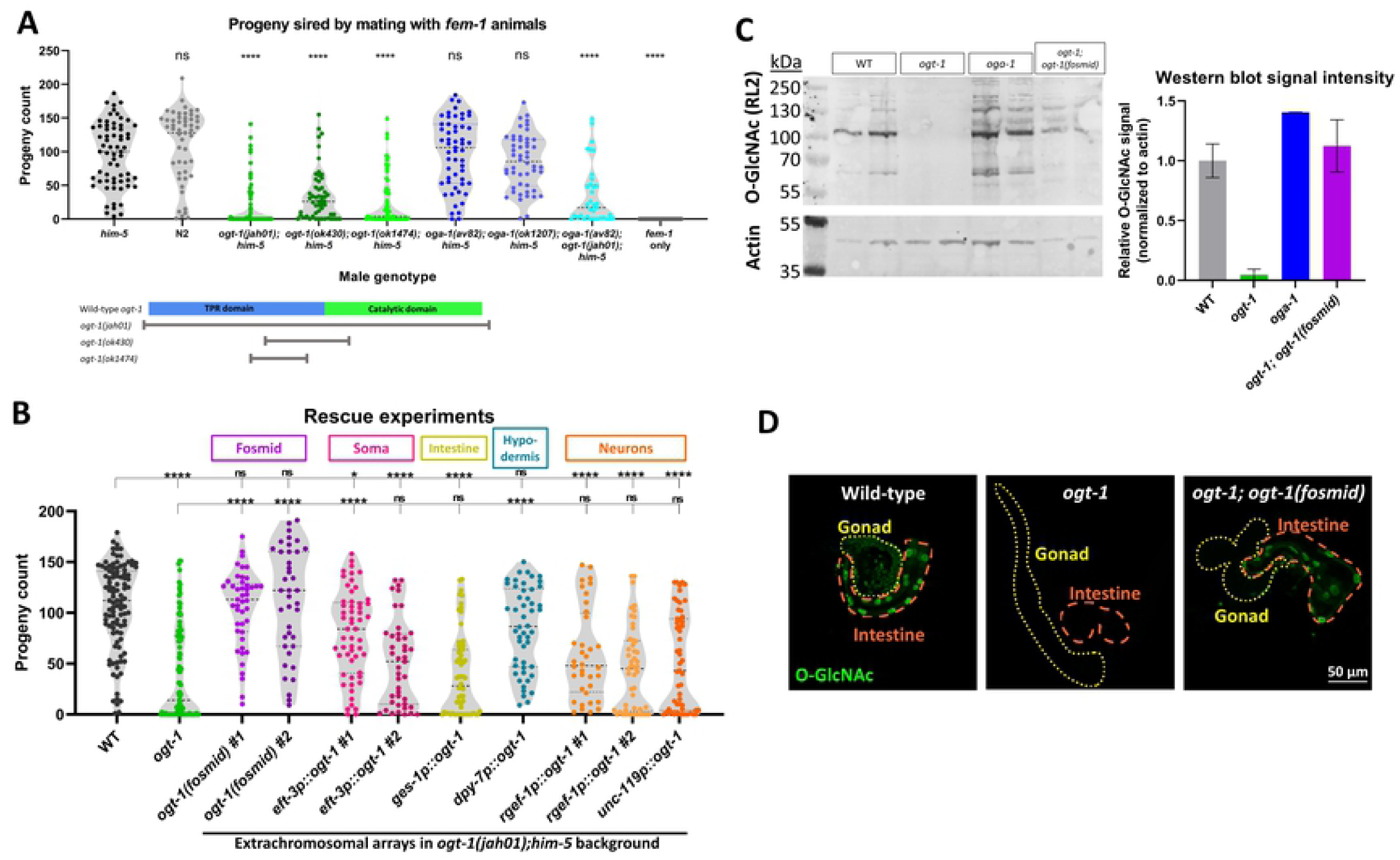
*ogt-1* mutations reduce male fertility. (A) 24-hour brood count from self-sterile *fem-1* worms mated with males of each specified genotype, shown with violin plots. Statistical comparisons to *him-5* by one-way ANOVA are shown. Below, a schematic of the three deletions in the context of the OGT-1 protein structure, with the tetracopeptide repeat (TPR) domain in blue and the catalytic domain in bright green. (B) 24-hour brood by mating with *fem-1*. All extrachromosomal array rescue lines are in the deletion background *ogt-1(jah01)*. Statistical comparisons by one-way ANOVA to WT and *ogt-1* are shown above. (C) Left, western blot of worm lysates demonstrates rescue of anti-*O-*GlcNAc antibody (RL2) signal in fosmid-rescued line. Sample for each lane is a pool of approximately 200 picked adults. Right, *O-*GlcNAc signal intensity per lane normalized to actin band, with WT set to 1. Error bars denote the range. See S1 Fig for uncropped blot. (D) Immunohistochemistry of dissected males shows *O-*GlcNAc staining is restored in somatic tissues (intestine, dashed orange line) of the fosmid-rescued line, but not in the germline (gonad, dotted yellow line). See S2 Fig for these images with additional channels: tubulin, DAPI, and differential interference contrast (DIC). All males in panels B-D are in the *him-5* background. * = p<0.05, **** = p<0.0001, ns = not significant.

Due to the ubiquitous expression of *ogt-1* throughout development [24, 32, 38] and the complexity of reproductive biology, the deficit in male fertility phenotype may originate from the many different tissues. To determine which tissue *ogt-1* expression is critical for normal fertility, we performed a series of tissue-specific rescue experiments. For these experiments, we used a CRISPR deletion of the whole gene, *ogt-1(jah01)*, as the background for genetic rescue. Through microinjection of a fosmid into this strain, we established two independent lines carrying a fosmid containing the *ogt-1* gene as an extrachromosomal array. Males with the fosmid had wild-type fertility when mated with *fem-1* worms (Fig 1B), demonstrating a robust rescue of the phenotype. To determine if functional OGT-1 was being produced by the fosmid-rescued lines, global *O-*GlcNAcylation was assessed by western blot. The blot showed *O-*GlcNAc levels were undetectable in *ogt-1(jah01)* worms (Fig 1C, lanes 3-4), and that the fosmid-rescued line restored global *O-*GlcNAcylation to wild-type levels (Fig 1C, lanes 7-8, S1 Fig), but not as high as the elevated levels detected from *oga-1(av82)* worms (Fig 1C, lanes 5-6). As an extrachromosomal array, expression of the *ogt-1* transgene was likely repressed in the germline [39], and we used antibody staining on the gonad and intestine of dissected males to test this. *O-*GlcNAc staining was present in both the intestine and gonad of wild-type worms, absent from both tissues in *ogt-1(jah01)* worms, and present in the intestine but not the gonad of the fosmid-carrying worms (Fig 1D). This suggests the fosmid enabled *ogt-1* expression in somatic tissues but not in the germline. With the fertility rescue, this indicates somatic expression of *ogt-1* was sufficient for wild-type fertility.

Because *ogt-1* is expressed globally [19, 24, 32], further experiments were conducted using extrachromosomal arrays to induce tissue-specific expression of *ogt-1* in a null mutant background to determine if any could rescue the fertility defect. Rescue plasmids were constructed using promoter fragments inserted directly upstream of the *ogt-1* cDNA containing two endogenous introns (see Materials and Methods for more detail). The same promoters were also cloned into a GFP plasmid and co-injected to ensure the promoters were expressing as expected. When pan-somatic-expressing *eft-3* promoter [40] was used to express *ogt-1* in the deletion background, males from two independent lines had higher average progeny count than deletion males (Fig 1B). This further confirmed somatic expression of *ogt-1* was sufficient to rescue male fertility, and that the *ogt-1* transgene was functional. Expression of *ogt-1* from the hypodermis-specific *dpy-7* promoter [41] was sufficient to rescue fertility (Fig 1B), suggesting the hypodermis is important to this phenotype. The intestinal promoter *ges-1* [41] and neuronal promoters *rgef-1* [42] and *unc-119* [43] did not rescue fertility (Fig 1B).

The male tail is composed of anatomical structures the animal uses in the mating process. Some of these structures, such as the sensory rays, are derived from hypodermal cells [44, 45], making male tail development an appealing hypothesis to explain the mating defect. Analysis of high-magnification images taken of the male tails revealed either a slight increase in developmental defects or no change from wild-type, depending upon allele (S3 Fig). The full-gene CRISPR deletion *ogt-1(jah01)* and smaller deletion *ogt-1(ok430)* both showed no significant elevation in defects. Twelve percent of *ogt-1(ok1474)* had developmental defects including missing rays or fusions of multiple rays, while these defects occurred in only 2% of wild-type males (S3 Fig). The defects we observed were primarily within the V6 rays (rays 4-6). Because this phenotype is specific to one allele and relatively mild (88% of males appear normal), it is unlikely to be a major contributor to *ogt-1* male infertility.

### *ogt-1* males transfer sperm less often

We next performed mating experiments to examine the cause of the fertility defect in greater detail. When a *C. elegans* male mates with a self-fertile hermaphrodite, the male sperm typically outcompete the hermaphrodite sperm, at nearly 100% efficiency [46]. To test if *ogt-1* male sperm can compete with hermaphrodite sperm, males were crossed with self-fertile *unc-39(ct74)* hermaphrodites. Self-progeny from *unc-39(ct74)* hermaphrodites had the uncoordinated (Unc) phenotype, while F1s resulting from male sperm were non-Unc. As expected, mating with wild-type males resulted in near 100% outcross progeny (Fig 2A), indicating successful sperm transfer and that the sperm was highly competitive. *ogt-1* males showed a bimodal distribution in percent outcross progeny, with many producing very high proportion of outcross progeny, and others showing zero or low percentages (Fig 2A). While *ogt-1* male sperm may have a competitive disadvantage, differences in mating behavior may also be important, as the slow-moving Unc hermaphrodites are easier to mate with than non-Unc worms.

**Fig 2.**
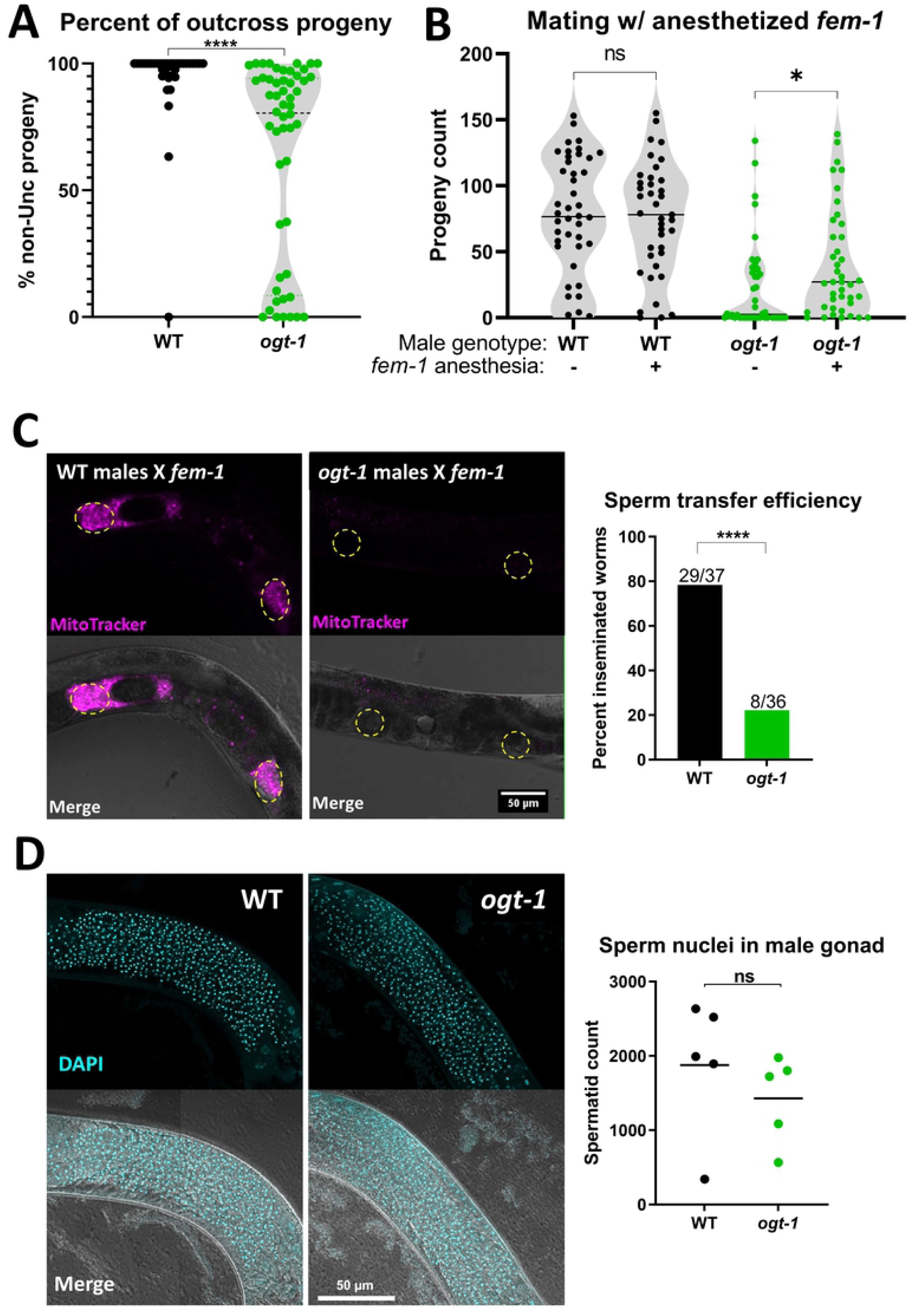
*ogt-1* males transfer sperm less often and have normal sperm count. (A) Sperm competition tested by mating males with *unc-39* hermaphrodites shows lower percent outcross progeny for *ogt-1* males, as assessed by t-test. (B) *ogt-1* males show higher progeny counts when mating with anesthetized *fem-1* worms, as assessed by t-test. (C) Representative images of unlabeled, unanesthetized *fem-1* animals after mating for one hour with MitoTracker-labeled males (magenta) of indicated genotype. Spermathecae are indicated with dashed yellow circles. Number of successful (any sperm in reproductive tract) versus unsuccessful sperm transfers (no detectable sperm) was compared between genotypes by Fisher’s exact test. (D) Representative images of DAPI-stained (cyan) virgin males, where spermatid nuclei appear as bright, compact puncta. Count of spermatid nuclei is unchanged between *ogt-1* and wild-type, as assessed by t-test. All males are in the *him-5* background. * = p<0.05, **** = p<0.0001, ns = not significant.

To directly test if crossing with *ogt-1* males with immobile mates results in more offspring, we tested mating with *fem-1* worms with and without anesthetic. Crossing with wild-type males resulted in many offspring whether or not their *fem-1* mates were anesthetized (Fig 2B). *ogt-1* males produced significantly more offspring when mating with anesthetized worms, but progeny counts were still lower than wild-type (Fig 2B). This improvement suggests initiation of mating was a significant barrier to reproduction for *ogt-1* males.

Sperm transfer can be directly assessed by tracking fluorescently labelled male-derived sperm within the reproductive tract of non-labelled mates. Unanesthetized *fem-1* worms were used as mates for consistency with the other experiments and to reduce the likelihood of ovulation which could displace male sperm. After one hour, 78% of *fem-1* worms mated with labelled wild-type males showed the presence of labeled sperm, while only 22% of those mated with *ogt-1* males did (Fig 2C). When sperm was present in *fem-1* worms mated with *ogt-1* males, there were fewer sperm, though the sperm did localize to the spermatheca as expected of sperm with normal motility (S4 Fig). By analyzing high-resolution z-stacks of DAPI stained virgin males, we were able to determine the number of spermatids is comparable between *ogt-1* and wild-type (Fig 2D). Taken together these data suggest behavioral differences may constitute a barrier to reproduction for *ogt-1* males.

### *ogt-1* males exhibit aberrant mating behavior

Due to *ogt-1* males’ better performance with slow-moving mates, we hypothesized they may have critical defects in mating behavior. The series of behaviors the adult male *C. elegans* must orchestrate in order to successfully mate are complex but well characterized [5]. A male must first locate a mate, then respond to physical contact by holding the ventral side of its tail against the hermaphrodite. The male will then search for the vulva by moving backwards along the length of the worm. When it reaches either the head or tail of the hermaphrodite, it will perform a tight turn to then search the other side for the vulva. Finally, the male must stop at the vulva, insert its spicules, and ejaculate into its mate.

Before the process of mating can begin, the male must actively seek out mates. The leaving assay was developed to measure the mate-searching behavior of *C. elegans* males. A lone adult male placed in a tiny lawn of OP50 will typically leave the food source to explore extensively due to its drive to find a mate (Fig 3A), while hermaphrodites will not leave the food source [3]. At the endpoint of the 24h assay, 62% of wild-type males had exhibited searching behavior, while only 38% of *ogt-1* males had done so (Fig 3B). To ensure this difference was not due to a movement defect, we analyzed video of *ogt-1* males using the wrMtrck ImageJ package [47] and found the average movement speed of males was equivalent to wild-type (Fig 3C). Thus, the leaving assay findings suggest *ogt-1* males have a reduced mating drive.

**Fig 3.**
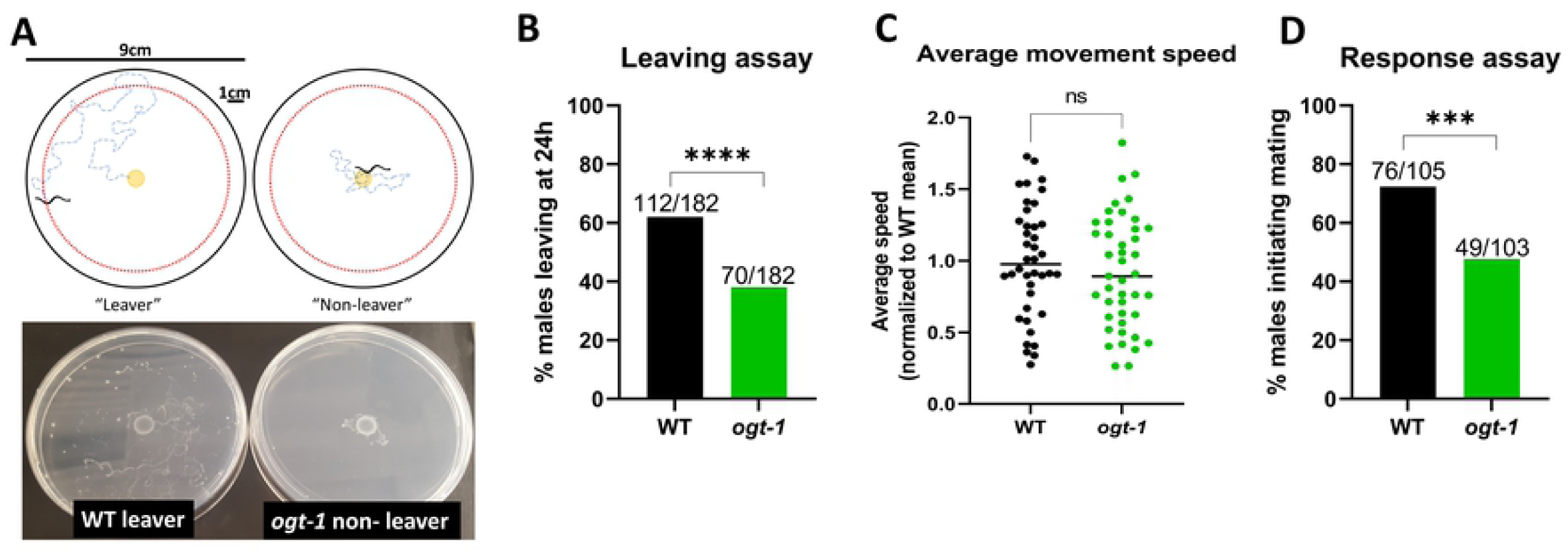
*ogt-1* males have reduced mating drive. (A) Schematic of the leaving assay: an individual males is placed in a small dot of OP50 (2μL) in the center of a 9cm plate and is scored as a “leaver” if it gets within 1cm of the edge of the plate over the course of the 24h assay. Below, images of representative plates illustrating the exploration of a wild-type leaver and an *ogt-1* non-leaver. For this image, plates were kept at room temperature after the assay to allow bacterial growth for easier visualization. (B) Leaving assay endpoint data (24h) shows *ogt-1* males are less likely to exhibit exploratory behavior associated with mating drive, as assessed by Fisher’s exact test. (C) Normalized average moving speed is no different between wild-type and *ogt-1* worms, as assessed by t-test. (D) Latency assay shows fewer *ogt-1* males initiate mating during a 6-minute assay, as assessed by t-test. All males are in the *him-5* background. *** = p<0.001, **** = p<0.0001, ns = not significant.

As the leaving assay assesses mating drive in the absence of mates, we next tested *ogt-1* males’ response to the presence mates. We scored mating initiation with a response assay adapted from prior studies of mating behavior [48, 49]. For this assay, initiation was defined as two seconds of contact between the ventral side of the male tail with its mate. Within a six-minute assay, wild-type males initiated mating 72% of the time, while *ogt-1* males did so only 48% of the time (Fig 3D). This significant difference supports the leaving assay results suggesting *ogt-1* males don’t seek mates as actively as wild-type.

To determine if *ogt-1* deletion caused detriments within further steps of the mating process, videos of males mating with anesthetized *fem-1* worms were analyzed. In the mating videos, less than half of males analyzed initiated mating. For those that do engage in mating, each male may initiate multiple times. Normalizing the number of initiation events to the total number of males tested yields an average of 0.87 initiations per wild-type male, and 0.62 initiations per *ogt-1* male (Table 1). Across the videos analyzed, *ogt-1* males engaged in 3-fold fewer total turns than control, normalized to the number of males tested (Table 1).

**Table 1.** Total and normalized instances of behaviors observed by genotype.

| Genotype | n | Initiations |  | Turns |  | LOVs |  |
| --- | --- | --- | --- | --- | --- | --- | --- |
|  |  | Total | per male | Total | per male | Total | per male |
| WT | 60 | 52 | 0.87 | 187 | 3.1 | 55 | 0.92 |
| *ogt-1(jah01)* | 115 | 71 | 0.62 | 120 | 1.0 | 41 | 0.36 |

n = total number of males assessed in video analysis
"per male" values represent the total instances of each behavior divided by n for each genotype.
LOV = location of vulva, instances of a male stopping at the vulva.

That *ogt-1* males turn less often while mating was also clear when assessed by comparing the number of turns of each individual male that engaged in mating (Fig 4A). *ogt-1* males also had poorer turn quality (as defined in [50]), with 12.5% of turns resulting in complete disconnection from their mate (“missed turns”), up from 1.6% in wild-type (Fig 4B). Successful turns that involved correction after loss of contact (“sloppy turns”) nearly tripled compared to wild-type (Fig 4B).

**Fig 4.**
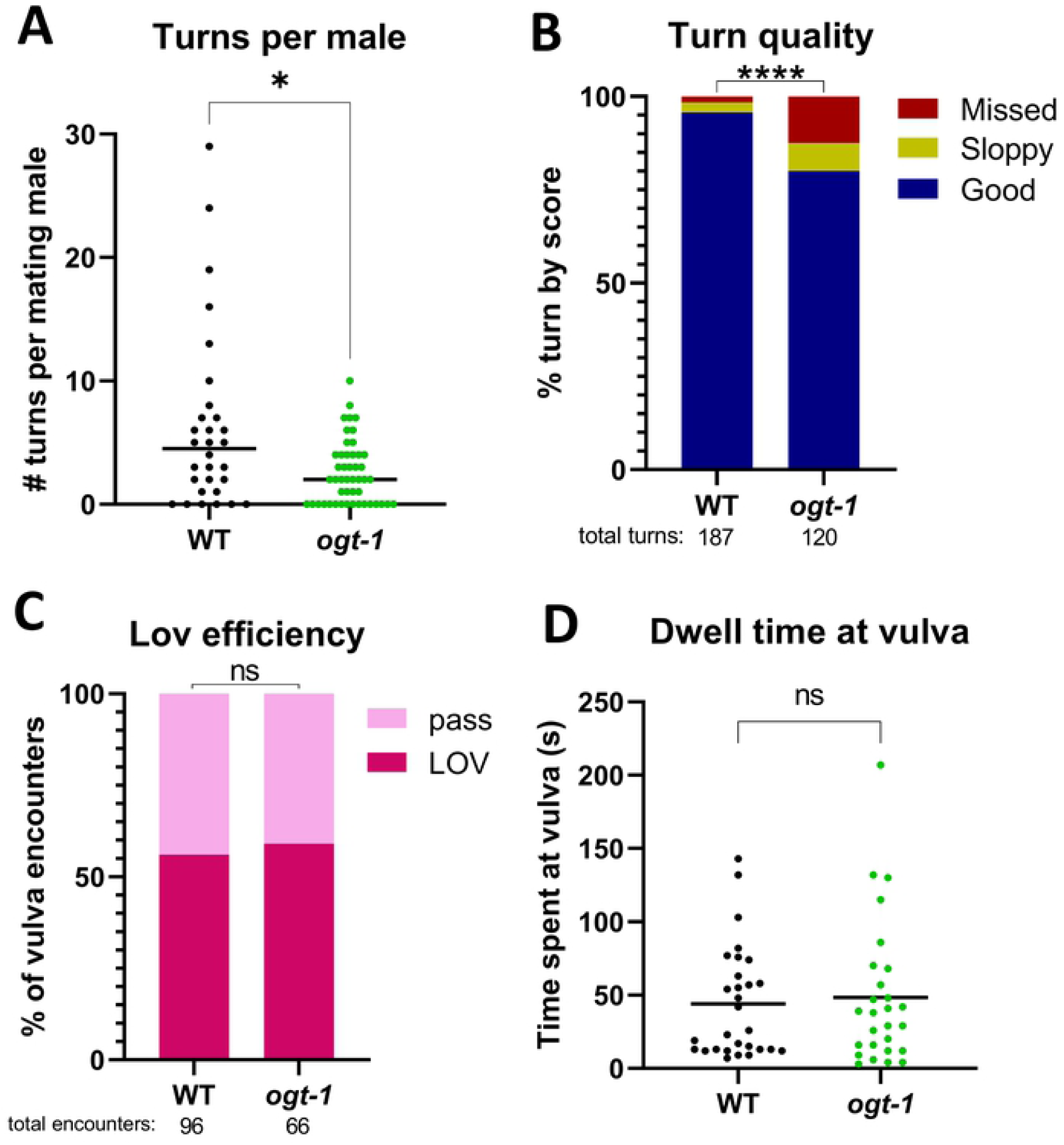
*ogt-1* males have reduced mating drive. (A) Over the course of mating, individual *ogt-1* males perform fewer turns on average, as assessed by t-test. Data includes only males which engaged in mating. (B) *ogt-1* males have a mildly higher incidence of sloppy turns (loss of contact, but successful turn) and of missed turns (complete loss of contact with mate), as assessed by Chi-square. (C) *ogt-1* males have normal vulval location efficiency, shown as the percent of vulva encounters in which the male showed successful location of vulva (LOV) versus the percent in which the male does not stop at the vulva (pass), as assessed by Fisher’s exact test. (D) *ogt-1* and wild-type males dwell at the vulva for similar lengths of time, as assessed by t-test. Dwell time is defined as the time from when the male stops at the vulva to the time the male leaves the vulva. All males are in the *him-5* background. * = p<0.05, **** = p<0.0001, ns = not significant.

*ogt-1* males had a 2.4-fold reduction in the total instances of successfully locating the vulva (Table 1). However, expressed as vulval location efficiency (successful vulval location events divided by total vulval encounters), *ogt-1* and wild-type were roughly equivalent (Fig 4C). This suggests that *ogt-1* males could properly identify the vulva but were less likely to encounter the vulva due to mating fewer times than wild-type, and searching their mates less thoroughly as suggested by the reduced total turns. Additionally, wild-type males and *ogt-1* males spent comparable amounts of time at the vulva after stopping (Fig 4D). These findings suggest the later steps of mating proceed normally in *ogt-1* males, supporting the idea that the mating drive deficit is the primary cause of the fertility defect.

### OGT-1 catalytic activity is dispensable for male fertility

Several recent studies have highlighted the importance of non-catalytic functions of OGT [14], including two that showed *C. elegans ogt-1* phenotypes did not require catalysis [24, 51]. To determine if OGT-1’s catalytic function was necessary for normal male fertility, we made use of two lines CRISPR-edited with H612A and K957M substitutions, constructed in the context of the fluorophore-tagged fusion *ogt-1::gfp* strain *ogt-1(dr84)* [24] (Fig 5A). These two sites correspond with the H498A and K842M sites in the human enzyme, which were among the lowest-activity OGT variants tested *in vitro* [52]. Western blot showed undetectable *O-*GlcNAc levels in both the H612A and K957M lines, comparable with *ogt-1* deletion (Fig 5B, lanes 6-7). Putting these alleles in the background of the *oga-1* CRISPR deletion *av82*, which cannot remove *O-*GlcNAc, we determined that H612A had very low levels of activity, as one light band was visible (Fig 5B, lane 8). *oga-1(av82);ogt-1[K957M]* did not show detectable signal (Fig 5B, lane 9), suggesting the K957M mutation more completely abrogates catalysis.

**Fig 5.**
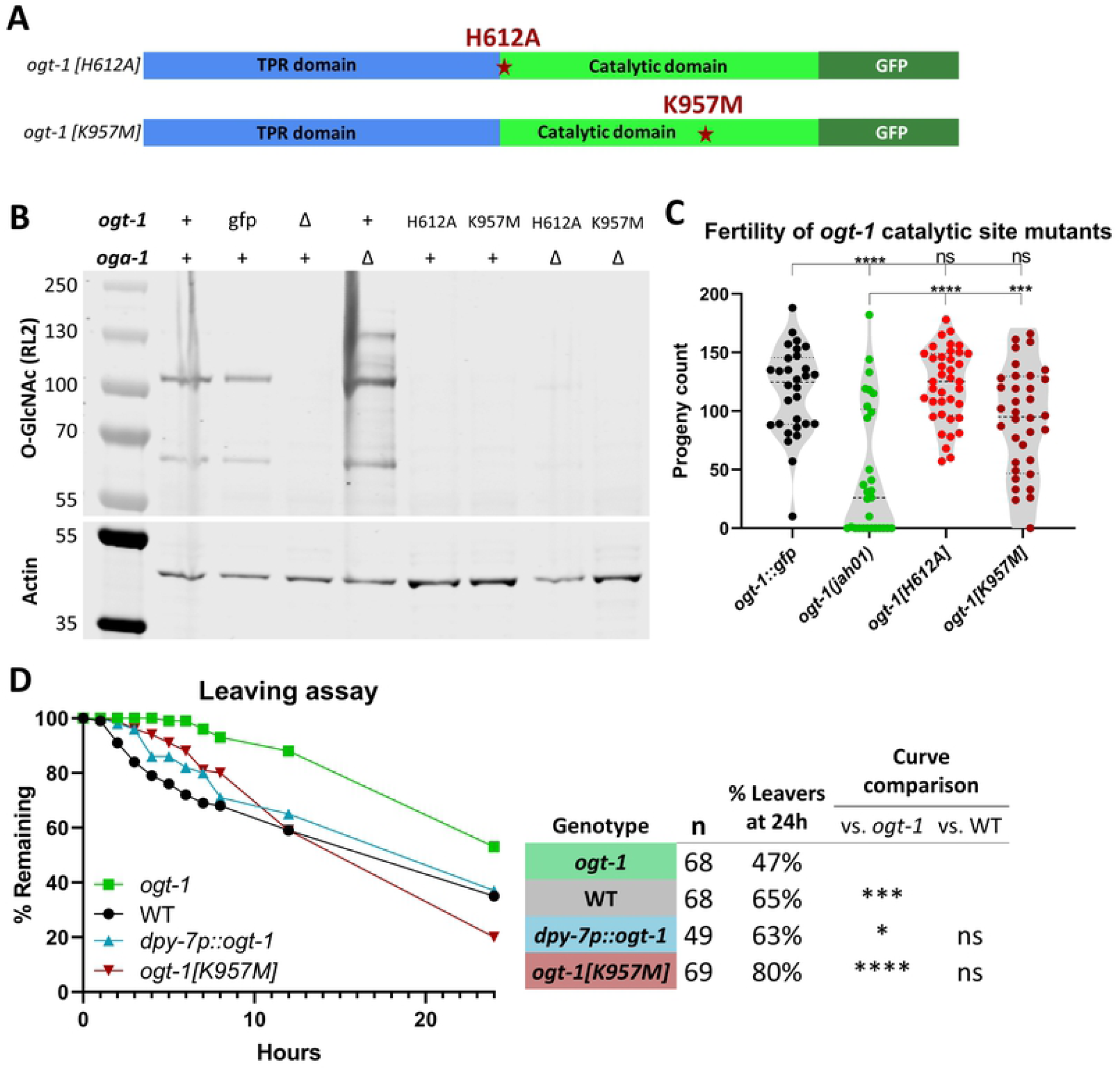
OGT-1 catalytic activity is not required for male fertility. (A) Schematic of OGT-1 protein showing the location of two substitutions within the context of the tetratricopeptide repeat (TPR) domain (blue), catalytic domain (bright green), and GFP tag (dark green). (B) Western blot of worm lysates shows undetectable *O-*GlcNAcylation in *ogt-1Δ (jah01)* and point mutant lines. When in the *oga-1Δ (av82)* background which increases global *O-*GlcNAc levels, *ogt-1[K957M]* has undetectable signal, and *ogt-1[H612A]* shows very faint signal. “gfp” is *ogt-1(dr84)[ogt-1::gfp]*, with similar *O-*GlcNAc signal to WT. (C) When mating with *fem-1, ogt-1* point mutant males have wild-type level of progeny, as assessed with one-way ANOVA comparisons to *ogt-1::gfp* and *ogt-1(jah01)*. (D) Leaving assay time course with datapoints at 0-18h, 12h, and 24h shows *ogt-1[K957M]* and *dpy-7p::ogt-1* males have similar mating drive to wild-type. Includes data from three replications of the experiment. Summary chart showing total males tested (n), the percent of leavers at the endpoint, and statistical comparison of curves by log-rank Mantel-Cox. All males are in the *him-5* background. * = p<0.05, *** = p<0. 001, **** = p<0.0001, ns = not significant.

Crossing with males from these missense mutants produced progeny counts comparable with the positive control *ogt-1::gfp*, and significantly higher than *ogt-1* deletion (Fig 5C). This demonstrates that male fertility requires OGT-1, but does not require *O-*GlcNAcylation, suggesting the protein is serving another role in this context.

As fertility can be impacted in a variety of different ways, we tested a catalytic-dead mutant and the hypodermal rescue line in the leaving assay to determine if their effects on progeny count were distinct from, or correlated with, mate-seeking. *ogt-1* males had the lowest rate of leaving behavior throughout the 24-hour assay, while the *dpy-7p::ogt-1* and *ogt-1[K957M]* males were similar to wild-type (Fig 5D). Curve comparison found significant differences between *ogt-1* and each of the other three tested lines. These results parallel the earlier findings that *dpy-7p::ogt-1* and *ogt-1[K957M]* males have wild-type levels of progeny when mating with *fem-1* mates. This correlation between behavior and progeny count phenotypes lends support to the hypothesis that the reduced progeny count of *ogt-1* males was caused by behavioral differences related to mating drive.

## Discussion

Here we have described a phenotype of *ogt-1* deletion that greatly impedes the fertility of males. Three independent alleles of *ogt-1* shared this phenotype, and expression of an *ogt-1* transgene either pan-somatically or specifically in the hypodermis was sufficient to rescue. *ogt-1* males transferred sperm less often and performed better with immobile mates. They were less likely to engage in mate-seeking behavior in the absence of mates, or to initiate mating when hermaphrodites are present. OGT-1 catalytic activity was not required for male fertility, as two lines with point mutations that abrogate *O-*GlcNAcylation had normal fertility. Both catalytic-dead and hypodermal rescue males displayed normal mate-seeking behavior, suggesting a non-catalytic OGT-1 function in the hypodermis is critical for male mating.

Adult *C. elegans* males, when isolated from mates, will spontaneously switch their behavioral paradigm from foraging to mate-seeking [3]. There is thought to be a certain probability that a male will begin mate search, which is captured by observing the rate of food-leaving over 24 hours using the leaving assay [3]. When tested in this way, *ogt-1* males were less likely to leave the food source, indicating a reduced mating drive as compared with wild-type males (Fig 3B). When mates are present, male food-leaving behavior is expectedly different, with males no longer engaging in the extensive search for a potential mate [3]. Using a different assay with an excess of mates present, we showed that *ogt-1* males in these conditions were less likely to initiate mating (Fig 3D). In addition to this, we observed *ogt-1* worms initiating mating less frequently in video analysis of mating behavior which used anesthetized *fem-1* worms (Table 1). Reduced mating initiation could be explained as a decreased response to contact with mates or could further reflect the reduced mating drive shown by the leaving assay. These findings may both be explained by a reduced mating drive for *ogt-1* males, or could reflect an aberrant response to contact which contributes to the low progeny counts in addition to mating drive. Lowering the barrier of entry to mating by using anesthetized or slow-moving mates improved the number of progeny produced by *ogt-1* males (Fig 2A-B), supporting the idea of a lowered but not completely ablated mating drive. Later in the mating process we also observed differences between *ogt-1* males and wild-type: fewer turns (Fig 4A, Table 1), a higher rate of poor turns (Fig 4B), and fewer successful stops at the vulva (Table 1). These results too may connect with the reduced mating drive, as fewer times engaging in mating should result in fewer instances of the downstream behaviors. Males with a reduced mating drive that do engage in mating may do so differently, searching for the vulva less actively or disengaging from mating before successful sperm transfer. Our results show a discrepancy between how often *ogt-1* males stop at the vulva, which was much lower than wild-type when normalized per male (Table 1) and equivalent to wild-type when normalized per encounter with the vulva (Fig 4C). This suggests *ogt-1* males were not deficient in their ability to detect the vulva but instead were encountering it less often, likely caused by aberrations in earlier steps of mating, especially mating initiation, turning defects, and mate search. Further highlighting the significance of mate-seeking, the *dpy-7::ogt-1* rescue line and the *ogt-1[K957M]* catalytic dead line were similar to wild-type in the leaving assay (Fig 5D), matching their results with the *fem-1* progeny count assay (Fig 1B, 5C). The behavioral assays did not show an effect as large as we observed for the sperm transfer assay (Fig 2C), which could suggest the behavioral differences in mate-seeking, response, and turning may add up to the greater effect seen downstream in the mating process. Together, our data show *ogt-1* males are different from wild-type in several mating behaviors, which may each contribute in part to low progeny counts.

A model of the conflicting drives between mate-seeking and feeding [4] would predict that since *ogt-1* males were seeking mating less, they therefore may be prioritizing feeding. This is in line with the role of OGT as a nutrient sensor, although this is complicated by OGT-1 catalytic activity being dispensable for male fertility (Fig 5C). OGT’s function as a nutrient sensor is typically thought to be contingent upon the concentration of UDP-GlcNAc which varies with cellular nutrient levels. Different nutrient conditions lead to OGT targeting different subsets of proteins for *O-*GlcNAc transfer [53]. Though these documented changes regard *O-*GlcNAc-modified proteins, they suggest more generally that OGT interacts with different proteins in high and low nutrient states. Interestingly, other TPR proteins are responsive to nutrients such as SSN6, a component of a glucose-responsive repressor complex in yeast [54]. In addition, expression of OGT is responsive to glucose and UDP-GlcNAc levels [53, 55, 56], though nutrient-dependent expression changes have not yet been tested in *C. elegans. ogt-1* deletion is known to change the metabolic profile of worms, with a 3-fold reduction in lipid stores, and 3-fold elevation in glycogen and trehalose stores [18]. These changes in metabolism may be interrelated with other *ogt-1* phenotypes such as shortened lifespan [21, 31], and reduced entry into diapause states including dauer [18, 29] and adult reproductive diapause [30]. Whether some or all these phenotypes depend on OGT-1 catalysis has not yet been determined, and merits further study. Starvation of males has been shown to reduce their probability of leaving their food source in the leaving assay [3, 57]. The metabolic changes seen in *ogt-1* mutants may induce a starvation-like response which may lead to some of the observed behavioral differences such as reduced mating drive.

We were surprised to find that the *dpy-7* promoter, which expresses predominantly in the hypodermis, was able to rescue the fertility phenotype (Fig 1B). The importance of *ogt-1* in the hypodermis was described in a recent study which found *ogt-1* is required to respond to hypertonic stress [24]. Hypodermis-specific expression (*dpy-7* promoter, as used in our study) of *ogt-1* was sufficient to restore normal response to high salt [24]. Also like our findings, catalytic-dead mutants were similar to wild-type in hypertonic stress response. Another study identified *ogt-1* in a mutagenesis screen for mutants which allowed transdifferentiation of adult hypodermal cells into neurons by overexpression of a neuronal transcription factor [32]. This indicates *ogt-1* helps maintain the hypodermal fate, a role it also plays in other tissues [33]. These findings further highlight the importance of *ogt-1* in the hypodermis and its non-catalytic functions.

The *C. elegans* hypodermis is an ectodermal epithelial tissue known for its role in producing the cuticle in conjunction with each molt [58]. It also regulates the growth of the organism [59, 60], stores nutrients [61], and forms important parts of certain anatomical structures such as the male tail [44], which appear to be largely normal in *ogt-1* males (S3 Fig). The hypodermis has also come to be understood as an important site of metabolism. A recent study of gene expression in different major tissues compared the gene expression profiles of adult *C. elegans* tissues with human tissues and interestingly found the *C. elegans* hypodermis to be most similar to the human liver [62]. This correlation is driven by the hypodermal expression of metabolic enzymes and regulators. An abnormal metabolic profile akin to starvation may shift behavioral prioritization towards food seeking, resulting in less engagement in mating. Among genes expressed in the hypodermis, lipid regulators are highly enriched [62]. Prior studies have shown *ogt-1* mutants are lipid-depleted [18, 30], which may connect with the role *ogt-1* plays in the hypodermis to affect fertility. Starvation is known to be one of the handful of perturbations which reduce mate-seeking behavior as measured by the leaving assay [3]. Prioritization of food over mate-seeking in starvation may involve a common pathway with *ogt-1*, and suggests other lipid regulators such as *nhr-49* would be interesting targets to investigate for their effect on mate-seeking behavior. Considering findings using mouse models, OGT may have an evolutionarily conserved role in regulating feeding behavior, as deletion of OGT in either the PVN neurons [63] or pancreatic α cells [64] induces overeating to the point of obesity.

Another pathway known to affect *C. elegans* mate-seeking is the insulin/IGF signaling pathway, which *ogt-1* is well-established as regulating [3, 4, 65]. Some phenotypes of *ogt-1* deletion parallel *daf-16/FOXO1* mutants such as shortened lifespan and reduced dauer entry [18, 21, 29, 31]. However, the effects diverge regarding mate-seeking, as *daf-16* worms show normal leaving [3] while *ogt-1* males are less likely to leave (Fig 3B, 5D). Additionally, the interaction of OGT with the insulin signaling pathway is thought to be mediated through *O-*GlcNAcylation of various pathway members [65], and the male fertility phenotype does not require catalysis (Fig 5C). Therefore, we do not favor the idea that this phenotype is mediated through the insulin/IGF signaling pathway. It has been proposed that starvation reduces mate-seeking behavior by reducing insulin signaling through *daf-2* [4], and our results suggest other pathways are likely involved and merit further study.

The *dpy-7* promoter is frequently used as a hypodermis-specific promoter in tissue-specific rescue experiments [60], but careful analysis of *dpy-7* expression has shown it turns on early in development in the P-lineage [66]. P-lineage cells produce many daughter cells fated to become adult hypodermis, yet some descendants differentiate into neurons [44]. Of particular interest are P10 and P11, which in the male produce several cells that form the hook sensillum [44]. This suggests an alternate interpretation of our data, whereby in the *dpy-7p::ogt-1* line, early expression of *ogt-1* in P10 and P11 may produce enough OGT-1 protein to be inherited by daughter cells which ultimately play a critical role in the hook sensillum of the adult male. However, the pan-neuronal promoters of *rgef-1* and *unc-119* did not rescue the fertility phenotype and should express in these neurons. Furthermore, males with hook sensillum cell ablations have difficulty finding the vulva (Lov phenotype) [45], while *ogt-1* males have Lov efficiency similar to wild-type (Fig 4C). Males with hook cell ablations were also noted to continue circling their mates, turning many more times than wild-type worms [45], while *ogt-1* males turn less often (Fig 4A, Table 1). Thus, the data from the genetic rescue experiments and behavioral analysis both suggest it is unlikely that *dpy-7p::ogt-1* rescues fertility due to expression in hook neuron precursors.

The male fertility phenotype we described here adds to a growing list of phenotypes which require the OGT-1 protein but not *O-*GlcNAcylation. This is the third non-catalytic role described in *C. elegans* [24, 51], and these are the only studies to have specifically tested the question of catalysis. The known functions of OGT other than *O-*GlcNAcylation include HCF-1 cleavage and other non-catalytic roles such as protein-protein interactions [14]. In invertebrates such as *C. elegans*, OGT does not cleave HCF-1 [67, 68]. OGT is required for development to adulthood in *Drosophila*, and studies have demonstrated early larval development requires *O-*GlcNAc, but the pupal lethality of OGT nulls is suppressed by variants with negligible *O-*GlcNAc transferase activity [69, 70].

Interestingly, the majority of patient alleles associated with OGT-XLID have occurred in the TPR domain [16], with only two variants found within the catalytic domain [71]. The TPR domain is not directly involved in *O-*GlcNAc transfer but is widely thought to be critical to OGT substrate selection and is the site of protein-protein interactions, several post-translational modifications, and dimerization [6, 72]. Of the OGT-XLID patient alleles, all but one retain the ability to transfer *O-*GlcNAc [73], though the enzyme kinetics of several variants are mildly reduced [15, 74]. The pathogenesis of OGT-XLID remains unclear and may be different for patients affected by different mutations. Several hypotheses have been proposed to connect the nature of these variants to the disorder, including reduced *O-* GlcNAcylation, lower OGA levels, HCF-1 misprocessing, or OGT destabilization [16]. With the evidence currently available, none of these four disease features appear to be shared between all OGT-XLID alleles [16]. A fifth hypothesis, alteration of the OGT interactome [16], is worth deeper investigation. Disruption of interactions between OGT and other proteins may prevent the *O-*GlcNAcylation of certain proteins which ultimately leads to the disorder, and it is also possible that these variants disrupt critical protein-protein interactions which may or may not involve glycosylation. In support of this conjecture, several regulatory interactions have been documented between OGT and other proteins that are likely wholly (Ataxin-10 [75]) or partially independent (p120-catenin [76, 77]) of OGT catalytic activity. We have argued that OGT-XLID pathogenesis may stem from disruption of OGT’s role regulating chromatin, DNA methylation, and gene expression [78]. Though OGT affects these processes through *O-*GlcNAc modification of various players in the respective pathways, non-catalytic functions likely compose an additional layer of regulation. The results we have reported here suggest there are additional non-catalytic roles of OGT to be discovered. Future studies in *C. elegans* and other organisms should test for divergence of function between the TPR and catalytic domains. As *C. elegans* tolerates loss of *O-* GlcNAcylation, this model system is uniquely suited to investigate non-catalytic roles of OGT and ultimately shed light on new pathways affected by OGT.

In conclusion, we have uncovered a previously undescribed defect in *C. elegans* male mating caused by *ogt-1* deletion, linked to a reduced mating drive. The phenotype was rescued by hypodermis-specific expression of an *ogt-1* transgene, suggesting the hypodermis plays a more important role in fertility than has been previously appreciated. Our experiments demonstrate that catalytic activity of OGT-1 is not required for fertility, adding to a growing list of non-catalytic roles of the enzyme. Further exploration of these functions will uncover new pathways regulated by OGT and build towards a better understanding of how disruption of OGT leads to disease.

## Materials and Methods

### Worm strains and maintenance

All worms were maintained at 20°C on NGM plates seeded with OP50 *E. coli* [79] unless otherwise noted. Strains were acquired through the Caenorhabditis Genetics Center at the University of Minnesota. Most strains were crossed into the *him-5(e1490)* background before they were used in experiments. Males from the following strains were used in this study: Bristol N2, CB1490 *him-5 (e1490) V*, DDK14 *ogt-1(jah01) III; him-5(e1490) V*, DDK11 *ogt-1(ok430) III; him-5(e1490) V*, DDK12 *ogt-1(ok1474) III; him-5(e1490) V*, DDK15 *him-5(e1490) V; oga-1(av82) X*, DDK13 *him-5(e1490) V; oga-1(ok1207) X*, DDK18 *ogt-1(jah01) III; him-5(e1490) V; oga-1(ok1207) X*. Hermaphrodites from two strains, BA17 *fem-1(hc17) IV*, and PD3165 *unc-39(ct73) V* were used in mating assays. The temperature-sensitive BA17 strain was maintained at the permissive temperature of 15°C [36, 80]. For rescue experiments, we established the following transgenic lines carrying extrachromosomal arrays: DDK16 *ogt-1(jah01) III; him-5(e1490) V; jahEx1[ogt-1(fosmid) + myo-2p::gfp]*, DDK19 *ogt-1(jah01) III; him-5(e1490) V; jahEx2[ogt-1(fosmid) + myo-2p::gfp]*, DDK23 *ogt-1(jah01) III; him-5(e1490) V; jahEx6[eft-3p::ogt-1 + eft-3p::gfp]*, DDK24 *ogt-1(jah01) III; him-5(e1490) V; jahEx7[eft-3p::ogt-1 + eft-3p::gfp]*, DDK25 *ogt-1(jah01) III; him-5(e1490) V; jahEx8[ges-1p::ogt-1 + ges-1p::gfp + myo-2p::gfp]*, DDK32 *ogt-1(jah01) III; him-5(e1490) V; jahEx11[dpy-7p::ogt-1 + dpy-7p::gfp + myo-2p::mCherry]*, DDK33 *ogt-1(jah01) III; him-5(e1490) V; jahEx12[rgef-1p::ogt-1 + rgef-1p::gfp + myo-2p::mCherry]*, DDK27 *ogt-1(jah01) III; him-5(e1490) V; jahEx9[rgef-1p::ogt-1 + rgef-1p::gfp + myo-2p::gfp]*, DDK31 *ogt-1(jah01) III; him-5(e1490) V; jahEx10[unc-119p::ogt-1 + unc-119p::gfp + myo-2p::mCherry]*. The *ogt-1::gfp* fusion and *ogt-1* point mutant lines were kindly gifted by the Lamitina lab, then crossed to *him-5(e1490)* to establish three strains used in this study: DDK26 *ogt-1(dr84)[ogt-1::GFP] III; him-5(e1490) V*, DDK28 *ogt-1(dr84[ogt-1::GFP] dr91[H612A]) III; him-5(e1490) V*, DDK29 *ogt-1(dr84[ogt-1::GFP] dr89[K957M]) III; him-5(e1490) V*. PCR was used for genotyping most strains; primers are listed in S6 Table.

### *fem-1* male mating assay

To test the ability of male worms to sire progeny, they were allowed to mate with *fem-1(hc17)* adults raised at 25°C which are effectively females, as they do not produce self-sperm [36, 80, 81]. For each cross, 15 males were plated with 5 *fem-1* animals on 35mm plate with a small lawn of OP50 and allowed to mate overnight (18h) at 20°C. After mating, each *fem-1* worm was transferred to a new plate and allowed to lay eggs over 24h, at which time the worm was removed. Progeny were subsequently counted 24-48h later. Males used in these assays were approximately 1-day adults. For strains carrying extrachromosomal arrays, only males positive for the fluorescent marker were used. One-way ANOVA was used to statistically compare progeny counts from each male genotype to wild type (*him-5*), or to *ogt-1*.

### *ogt-1* rescue experiments

A 30kb genomic fragment fosmid (WRM0635dF05) containing wild-type *ogt-1* was injected into *ogt-1(jah01);him-5* hermaphrodites along with a fluorescent marker (*myo-2p::gfp*) to create two independent rescue lines.

In order to test tissue-specific expression of *ogt-1* on the fertility phenotype, several plasmids with different promoters were constructed. To insert the *ogt-1* gene into a plasmid, the longer isoform of the gene (K04G7.3a.1) and its 3’UTR (614bp) were amplified from cDNA. Primers added an XmaI restriction site before the start codon, and a SpeI restriction site following a putative polyadenylation site downstream of the 3’UTR. This PCR product was cloned into a pBlueScript plasmid backbone. As introns have been reported to enhance transgene expression [82], the endogenous introns 5 and 6 (each approximately 50bp) were inserted into the gene by amplifying a portion of the gene from genomic DNA and using restriction sites for StyI and BlpI within exonic sequence. This intron-containing plasmid was used for subsequent cloning to add various promoters, which was done by ligating in promoter fragments amplified from genomic DNA using primers to add a SalI site upstream and a XmaI site downstream. These same promoter fragments (with one added nucleotide to keep GFP in frame) were also cloned into the promoterless GFP vector pPD95.67 (originally provided by Dr. Andrew Fire) to co*-*inject with the *ogt-1* rescue plasmids. See S6 Table for the primers used for genotyping and cloning.

Lines with tissue-specific *ogt-1* expression were established by microinjection of these plasmids into *ogt-1(jah01);him-5* hermaphrodites. Each injection mix contained the *ogt-1* rescue plasmid, the GFP expression plasmid with the same promoter, a fluorescent marker such as *myo-2::mCherry*, and empty pBlueScript to dilute.

### Western blot

Samples used for western blotting were either approximately 200 picked adults (Fig 1C, S1 Fig), or washed off one 6cm plate with many adults (Fig 5B, S6 Fig). Samples were then directly boiled in SDS loading dye (Quality Biological) w/ freshly added 2-mercaptoethanol to 2%. Samples were run on polyacrylamide gel in MOPS, then transferred to nitrocellulose at 100V for 100 minutes, or dry transferred using the iBlot 2 system. Primary antibodies targeting actin (Abcam ab8227) and *O-*GlcNAc (RL2, Thermo MA1-072) were used, followed by Odyssey secondary antibodies. The stained membrane was imaged by Odyssey.

### Immunohistochemistry

Worms were put in buffer on poly-lysine treated slides and dissected using two needles to remove the head. Immediately following dissection, a coverslip was applied, and the slide was immersed in liquid nitrogen. The coverslip was then flicked off, and the slides were immersed in methanol for fixation. The RL2 antibody (Thermo MA1-072) was used to target *O-*GlcNAc, and tubulin was visualized with an antibody targeting TBB-2 kindly gifted by Dr. Kevin O’Connell. Fluorescent AlexaFluor secondary antibodies (Invitrogen) were used for visualization. Slides were mounted using Vectashield with DAPI (Vector labs) and imaged with a 20X objective on a Zeiss LSM 700 confocal microscope.

### Live worm imaging of male tail

On glass microscope slides, 2% or 5% agarose was used to make pads. Young adult males were picked into M9 containing anesthetic on the agar pads, then a coverslip was applied, and the worms were immediately imaged with a 63X oil objective on a Zeiss LSM 700. Male tails were scored as deviant if any of the V6 rays were missing or fused.

### Sperm competition

Since hermaphrodites produce their own sperm, *C. elegans* male sperm have a competitive advantage to preferentially fertilize oocytes [83]. To assess the competitiveness of *ogt-1* male sperm [84], males were crossed with self-fertile hermaphrodites with a scorable phenotype: the uncoordinated phenotype (Unc) of *unc-39(ct73)* homozygotes. Crosses were performed overnight at 20°C with 15 males and 5 *unc-39(ct73)* hermaphrodites on a 35mm plate with a small OP50 lawn. The 24h brood of each hermaphrodite was then allowed 24-72h to develop to make the Unc phenotype easier to discern. The number of Unc and non-Unc progeny was counted and reported as the percent of non-Unc out of total progeny and compared by t-test. As *unc-39(ct73)* hermaphrodites used in this assay had relatively high rates of death, the data reported represents only those that were still alive and healthy after the 24h period of egg laying.

### Sperm transfer

As has been previously described [85], MitoTracker was used to stain live males and thus fluorescently label their sperm. Males were stained with 10μM MitoTracker Red (Invitrogen) in a watch glass for two hours, then put on a normal NGM plate to recover overnight. These males were then plated with unlabeled mates. We chose to use *fem-1* females for consistency with other experiments and to reduce the chance that ovulation would displace sperm. *fem-1* worms were then separated from the males and allowed to recover for one hour before mounting on 2% agar pads and imaged with a 20X objective on a Zeiss LSM 700. For this experiment, sperm transfer success was defined as any MitoTracker-labeled sperm present in the reproductive tract of the *fem-1* animal. The number of instances of mating success and failure by each male genotype were compared using Fisher’s exact test.

### DAPI staining of virgin males for sperm count

To ensure males are virgin prior to assessment of sperm count, synchronized males at the L4 stage were picked onto a plate and allowed to develop into adults overnight. These adult virgin males were then fixed dehydrated by applying successively higher ethanol concentrations, then fixed with acetone and stained with 20μg/μL DAPI in PBS. Each slide was imaged with a 63X oil objective on a Zeiss LSM 700 as a z-stack with 0.2um steps. Analysis of these images was performed with Imaris software using a published method for counting germline nuclei [86], with parameters adjusted for the size of spermatid nuclei rather than hermaphrodite germ cell nuclei.

### Mating behavior

The leaving assay [3] was used to score mate-searching behavior. 9cm plates were poured thin with 10mL of NGM, seeded the next day with 2μL of OP50 liquid culture, then used in the assay the following day. A single male was placed in the tiny OP50 lawn and monitored over 24h to see if it has engaged in exploratory behavior by looking at the tracks left by the male on the plate by eye. Plates were checked every hour for the first 8 hours, then at the 12h and 24h timepoints. A male is considered a “leaver” if its tracks reach 1cm or closer to the edge of the plate at some point over the course of the assay. The number of leavers and non-leavers for each male genotype at the endpoint were compared using Fisher’s exact test. Curve comparison between multiple genotypes was performed with the log-rank Mantel-Cox test.

Mating initiation was scored by the response to contact assay [48, 49]. *fem-1* worms were plated on a 35mm NGM plate with a small OP50 lawn and allowed an hour to condition the plate prior to the start of the assay. For each assay, 5 males (either WT or *ogt-1*) were added to the center of the OP50 lawn and allowed up to 6 minutes to initiate mating, here defined as 2 seconds of contact between the ventral side of the male tail and the mate. Males that initiated mating were removed from the plate during the assay. The proportion of mating initiation and failure to initiate between genotypes was compared using Fisher’s exact test.

The average movement speed of individual males was determined by recording and analyzing videos of adult males moving freely on an unseeded NGM plate. To capture video, an Olympus DP74 camera was used with an Olympus SZX16 dissecting microscope. The videos were analyzed using the wrMtrck software package for ImageJ [47], which tracked the movement of each individual male throughout the duration of each video.

Video analysis was used to assess turns, location of vulva (Lov), and time spent at the vulva. Videos were recorded with the Olympus DP74 camera on an Olympus SZX16 dissecting microscope over five- or ten-minute intervals. Several *fem-1* worms were anesthetized with 0.1% tricaine and 0.01% tetramisole just prior to the experiment and were distributed evenly on the OP50 lawn throughout the viewing area. An equal number of males were then added to the plate and the recording started. Videos taken on the same day alternated between wild-type and *ogt-1* male genotypes, using the same plate and *fem-1* animals for consistency. Initiation was defined as contact between the male tail and its mate and the start of backward locomotion. Turns were assessed as “good”, “sloppy”, or “missed” as has been defined in prior studies of turning behavior [50]. The total number of turns in each category for control and *ogt-1* were compared by Chi-square analysis. The ability of a male to locate the vulva (location of vulva or LOV) was scored by the ratio between the number of times the male stopped at the vulva to the total number of times the tail passed over the vulva without stopping [45, 87]. Dwell time at the vulva was defined as the interval from when the male located the vulva to the time the male disengaged from the vulva. In cases where the video ended before the male disconnects, or where another male knocks the male off, these dwell time values were excluded from the analysis. We also report the total number of behaviors (initiations, turns, vulval locations) per genotype, normalized to the total number of males analyzed in the videos.

### Statistical analysis

Comparisons between groups were performed with t-test for two groups, or one-way ANOVA for three or more groups, using GraphPad Prism 9 (GraphPad Software, Inc., La Jolla, CA). When comparing binary categorical data, Fischer’s exact test was used. Chi-square was used for turn quality analysis (Fig 4B), which included three categories. For Fig 5D, curve comparison was performed with the log-rank Mantel-Cox test.

Funding: This work was supported by NIDDK intramural funds (NIH). The funders had no role in study design, data collection and analysis, decision to publish, or preparation of the manuscript.

## Acknowledgements

We thank the members of the Hanover lab, Dr. Andy Golden, Dr. Xiaofei Bai, and the other members of the Golden lab, Dr. Harold Smith (NIDDK Genomics Core), and the Baltimore Worm Club for helpful discussions, technical assistance, reagents, and strains. We thank Dr. Kevin O’Connell for his help with immunohistochemistry and for sharing his anti-TBB-2 antibody. We thank the Lamitina lab for sharing strains.

## Author contributions

Conceived and designed the experiments: DK TF MWK JAH. Performed the experiments: DK. Analyzed the data: DK MD. Original draft preparation: DK. Supervision and critical reading of the manuscript: TF MWK JAH.

---

**S1 Fig. Uncropped western blot with fosmid rescue line from Fig 1C**

Uncropped western blot. Left: *O-*GlcNAc (RL2 antibody), right: actin antibody. All genotypes have *him-5* in background.

**S2 Fig. Immunohistochemistry of *ogt-1*(fosmid) line, with all channels shown**

Immunohistochemistry of dissected males, as shown in Fig 1D, including *O-*GlcNAc (RL2 antibody, in green), tubulin (anti-TBB-2 antibody, in red), DAPI (in blue), and all channels merged with differential interference contrast (DIC, greyscale).

**S3 Fig. One *ogt-1* allele has increased incidence of ray development defects**

(A) Representative DIC image of a normal male tail for wild-type control and an example of fused rays (arrow) in the *ogt-1(ok1474)* line. (B) Percentage of males with normal V6 ray morphology, with total number of observed normal and total males of each genotype. Defects include ray fusions and missing rays. Ratio of normal to deviant males compared to WT with Fisher’s exact test. ** = p<0.01, all others not significant. All males are in the *him-5* background.

**S4 Fig. *ogt-1* males transfer fewer sperm, sperm localizes to spermatheca**

Three representative DIC (greyscale) images of *fem-1* animals after successful sperm transfer from MitoTracker-labelled (magenta) males of indicated genotypes. These images include only successful sperm transfer, excluding those with no sperm in their reproductive tract. Success rate (shown in Fig 2C) included with genotype above images.

**S5 Fig. *ogt-1* males are less likely to leave food**

Time course of the leaving assay, of which the 24h timepoint is shown in Fig 3B. Each point represents the percent remaining at each timepoint, with data summed from six replications of the leaving assay. Three replications of the experiment are not included in this figure, as they did not include all time points (all nine experiments are included in Fig 3B). Data was collected for each hour from 0-8, and at 24h. All worms are males in the *him-5* background. Curve comparison done between genotypes with log-rank Mantel-Cox test. **** = p<0.0001

**S6 Fig. Uncropped western blot with *ogt-1* point mutant lines from Fig 5B**

Left: *O-*GlcNAc (RL2 antibody), right: actin antibody. Genotypes of strains given above with regard to the *ogt-1* and *oga-1* genes. gfp = *ogt-1(dr84)*, Δ = deletion (*ogt-1(jah01)* or *oga-1(av82)*), H612A = *ogt-1(dr91)*, K957M = *ogt-1(dr89)*. All strains are in the *him-5* background.

